# Optimizing Coarse-Grained Models for Large-Scale Membrane Protein Simulation

**DOI:** 10.1101/2024.05.13.594009

**Authors:** Chen Yun Wen, Yun Lyna Luo, Jesper J. Madsen

## Abstract

Coarse-grained (CG) models have been developed for studying membrane proteins at physiologically relevant scales. Such methods, including popular CG lipid models, exhibit stability and efficiency at moderate scales, but they can become impractical or even unusable beyond a critical size due to various technical issues. Here, we report that these scale-dependent issues can arise from progressively slower relaxation dynamics and become confounded by unforeseen instabilities observed only at larger scales. To address these issues, we systemically optimized a 4-site solvent-free CG lipid model that is suitable for conducting micron-scale molecular dynamics simulations of membrane proteins under various membrane properties. We applied this lipid model to explore the long-range membrane deformation induced by a large mechanosensitive ion channel, PIEZO. We show that the optimized CG models are powerful in elucidating the structural and dynamic interplay between PIEZO and the membrane. Furthermore, we anticipate that our methodological insights can prove useful for resolving issues stemming from scale-dependent limitations of similar CG methodologies.

## Introduction

Cell membranes are essential in biology. They are responsible for separating the cell from its surroundings and enabling structural deformations in response to mechanical stress.^1^ The creation and detection of cell membrane deformation by membrane-embedded proteins are critical for probing the physical environment surrounding cells and transducing various mechanical stimulations into electrochemical signals.^2^ Computational simulation techniques such as molecular dynamics (MD) are increasingly trustworthy and useful in studying membrane proteins, in part due to their high space–time resolution. ^3–6^ However, using MD simulations at atomic resolution or near-atomic resolution (e.g., united atom^7,8^ or MARTINI models^9^) to simulate cellular processes involving membrane proteins at micron scale that occur extremely slowly (compared to the integration time step, which is typically a couple of femtoseconds) has not generally been feasible. Highly coarse-grained (CG) approaches, where significantly more than 4 heavy atoms and associated hydrogens maps into a single CG bead (approximately the MARTINI model resolution^9^), is one way to overcome limitations of scale when the atomistic detail is less relevant to the question under investigation.

Here, we first tested the limit of a previously developed CG lipid model that assembles into a bilayer membrane, the Grime lipid model (Fig. 1),^10,11^ for simulating a submicron scale membrane patch. While there are many choices of similar CG lipid models, ^12–26^ the advantage of the Grime lipid is its solvent-free phenomenological character that has proven efficient and tunable.^10^ The Grime model has been successfully applied to such diverse applications as protein scaffolding,^27–31^ transmembrane proteins,^32^ lipid droplets,^33–35^ lipid nanocarriers,^36^ and viral particles.^37–39^ Unfortunately, we observed that above a critical size, practical limitations in the use of the CG lipid model for describing bilayer membranes can become overwhelming. Specifically, the scale-dependent issues can come in the form of (transient or persistent) pores, which were not present in smaller-scale simulations using identical parameters, and excessive membrane undulations that distort and buckle the planar bilayer until complete collapse. Our observations appear different in nature to other issues arising from poor technical choices, such as updating the neighbor list too infrequently, which also can give rise to unphysical distortions in bilayer simulations.^40^ We demonstrate that by optimizing the lipid parameters, the problems of pores and excessive membrane undulation can be eliminated. E.g., adjusting the molecular ratio of lipids (by reducing the size of the head bead) resolves poration issues. However, in order to simultaneously solve the issue with sinusoisal distortions caused by non-vanishing lateral press or spontaneous curvature, modifications to both head and interface beads were found necessary.

**Figure 1:**
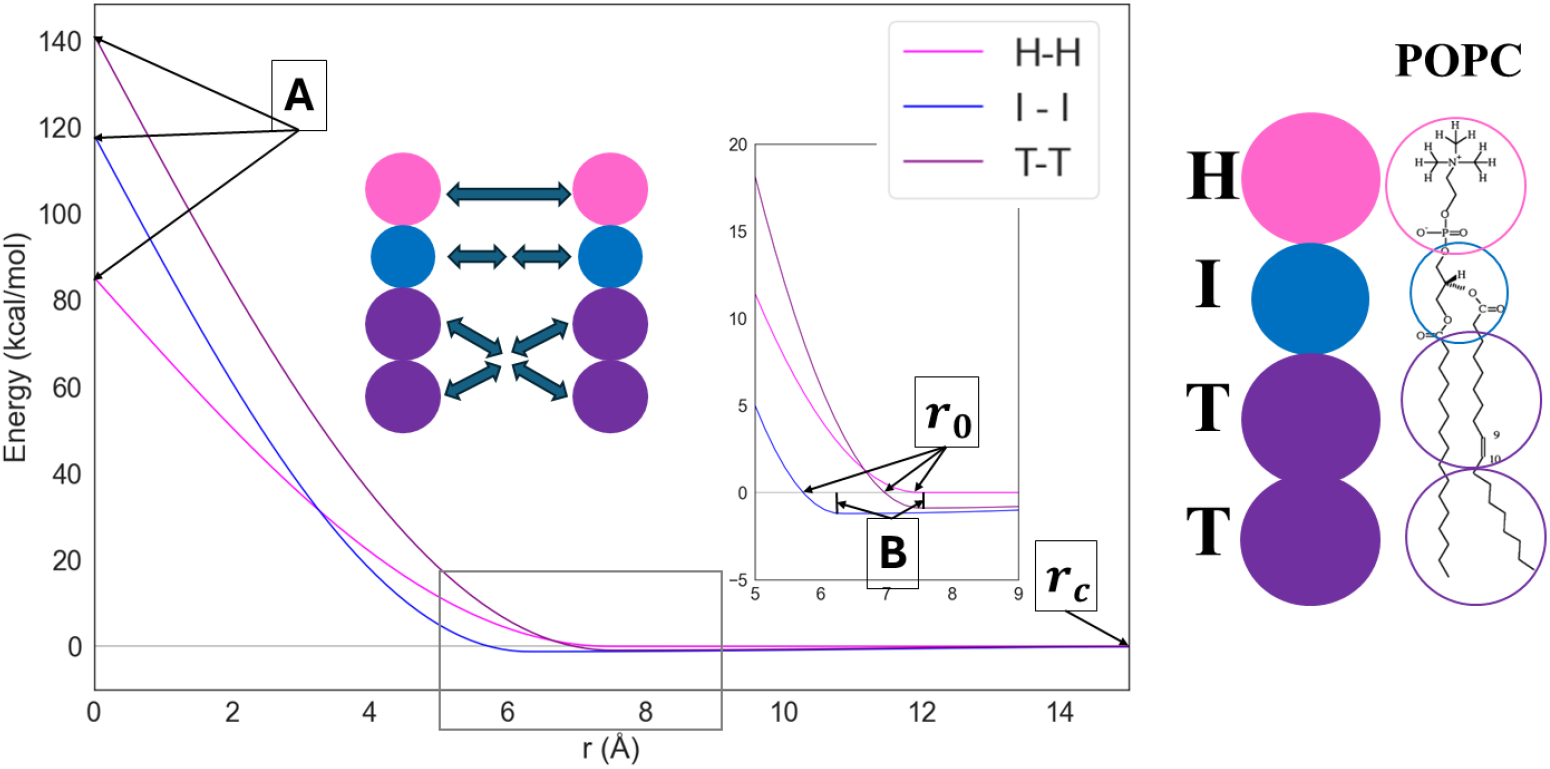
Left: Pairwise potential of the intermolecular interaction in the CG lipid model. The head bead interaction, *H* − *H*, is all-repulsive (*B* = 0), whereas an attractive cohesion is in effect for the *I* − *I* and the *T* − *T* interaction. Cross-interactions between different bead types are all-repulsive (*B* = 0). Example parameter values for *A, B, r*_0_, *r*_*c*_ are used. For actual parameter used in simulations, refer to the Methods section. Right: Schematic representation of the 4-site coarse-grained (CG) lipid model. Head (“H”), interface (“I”) and tail (“T”) type beads are shown as magenta, blue, and purple spheres, respectively. An approximate atomic mapping into corresponding CG beads is indicated for the prototypical lipid 1-palmitoyl-2-oleoyl-glycero-3-phosphocholine (POPC).

We next tested whether this newly optimized lipid model can capture the long-range membrane deformation by one of the largest plasma membrane proteins (totaling ∼ 8000 residues), called PIEZO.^41,42^ PIEZO1 and PIEZO2 are the main mechanosensitive ion channels in vetebrates. Their homotrimeric structures consist of a central ion permeating pore region and three large transmembrane mechanosensory domains (36 transmembrane helices each), known as “arms” or “blades”. In the absence of mechanical forces, the arms are curved, giving the protein an inverted dome shape (dome surface area ∼ 700*nm*^2^).^43^ This dome-shaped conformation imposes a curvature to the membrane around the channel, known as PIEZO membrane “footprint”.^44^ Due to the membrane bending rigidity, PIEZO’s footprint is predicted to extend measurably beyond the protein structure and decay gradually. Using a membrane patch of 115x115 *nm*^2^, we simulated PIEZO’s footprint under various membrane conditions using an elastic network model (ENM) of PIEZO. The ENM protein model is remarkably effective while being very simple, accurately capturing Gaussian fluctuations,^45–47^ even occasionally for proteins with multiple conformational states.^48^ For symmetric transmembrane proteins like PIEZO, it is advantageous to explicitly ensure that the CG mapping satisfies the structural symmetries.^49–51^ Generalizing and extending CG models for proteins opens promising avenues, ^52–54^ especially for PIEZO, but this is beyond scope here.

In summary, we demonstrate that a significant extension to the useful range of our CG simulation setup can be achieved by optimizing the lipid parameters, making it accessible to study long-ranged mechanical coupling between protein and lipids. Simulation of the optimized model predicts that increases in membrane tension or membrane bending rigidity lead to a reduction in PIEZO’s membrane footprint. The CG model encompasses proteins that are fully embedded within the membrane, which contrasts with the majority of prior (CG)MD studies that focused on membrane curvature effects (sensing or generation) resulting from proteins bound to membrane surfaces.

## Models and methods

### Model parameterization

#### Lipid model: 4-bead Grime lipid

The potential energy function of the Grime CG lipid model is given by^10^

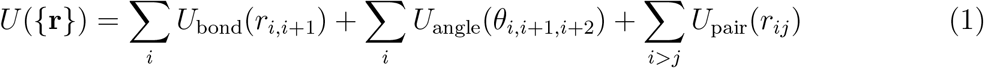

with

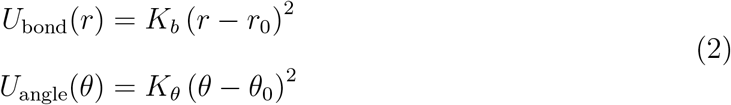

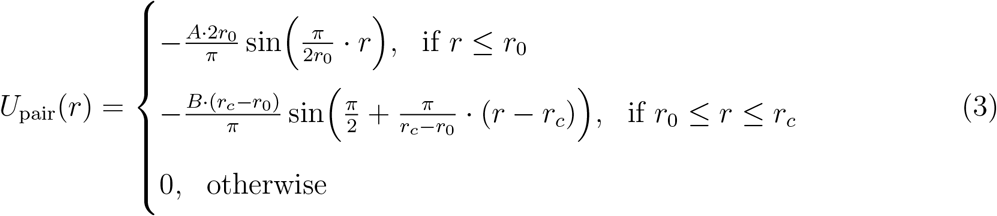

where *K*_*b*_ = 25*k*_*B*_*T Å*^*−*2^ is the bond force constant, *K*_*θ*_(*k*_*B*_*T* · rad^*−*2^) is the angle potential constant that controls the membrane bending rigidity (i.e., BEND), *r*_0_ is the bead size, and *r*_*c*_ is the interaction cutoff distance. In the 4-bead variant, each lipid molecule consists of a head bead (“H”), an interface bead (“I”), and two tail beads (“T”) (Fig. 1). The mass of each CG bead was set to be equal by evenly distributing the the mass of one 1-palmitoyl-2-oleoylglycero-3-phosphocholine (POPC) lipid over the four beads. The reference area-per-lipid for POPC is 64.3 ± 1Å^2^(experiment), ^55^ which is also consistent with all-atom MD simulations (e.g., using the CHARMM36 force field^56^).^57,58^

Using the original proposed set of parameters (*A* = 50*k*_*B*_*T, B* = 0.311*k*_*B*_*T, B*_*cross−terms*_ = 0, *r*_0_(*H* −*H, I* −*I, T* −*T*) = {0.75, 1, 1}*𝓁* = 7.5*Å, r*_*c*_ = 2*r*_0_),^10^ the resulting bilayer membrane for a system of 20*k* lipids can be controlled via a single parameter, the angle potential coefficient *K*_*θ*_ (or the equivalent, unit-less equivalent, BEND/(*k*_*B*_*T* · rad^*−*2^)), to yield bending stiffness ranging from a few to ∼ 100*k*_*B*_*T* without any noticeable issues. The optimized set of parameters is given by modification from the original set as follows. The bead sizes are *r*_0_(*H* − *H, I* − *I, T* − *T*) = {1, 0.84, 1}*𝓁* and the range of the *I* − *I* cohesion is increased by 10%, *r*_*c*_(*I* − *I*) = 1.1 · *r*_*c*_.

### Protein model: CG ENM

PIEZO is modeled as an elastic network model where harmonic bonds connect CG beads. To match the lipid resolution, the average resolution of the PIEZO model is ∼ 3 amino acids per CG bead. The beads are typed so as to match the character of the bilayer membrane (traversing the membrane, the bilayer has the types H-I-T-T T-T-I-H). Non-membrane-interacting bead types of PIEZO (e.g., loops or the “Cap” domain and the ‘beam’) are typed as head bead types, which are all-repulsive.

Since the CG beads are connected by the harmonic bonds, a cutoff of 16.25*Å* was determined to be sufficient for maintaining protein stability and flexibility. The equilibrium distances for the harmonic bonds were binned using bins of 0.25*Å* to reduce the required number of bond types. All harmonic bonds were assumed the same force constants, and this value was determined to be 2 (*kcal/mol*)*/Å* by comparing root-mean-squared fluctuation to a reference all-atom MD simulation (Fig. 2).

**Figure 2:**
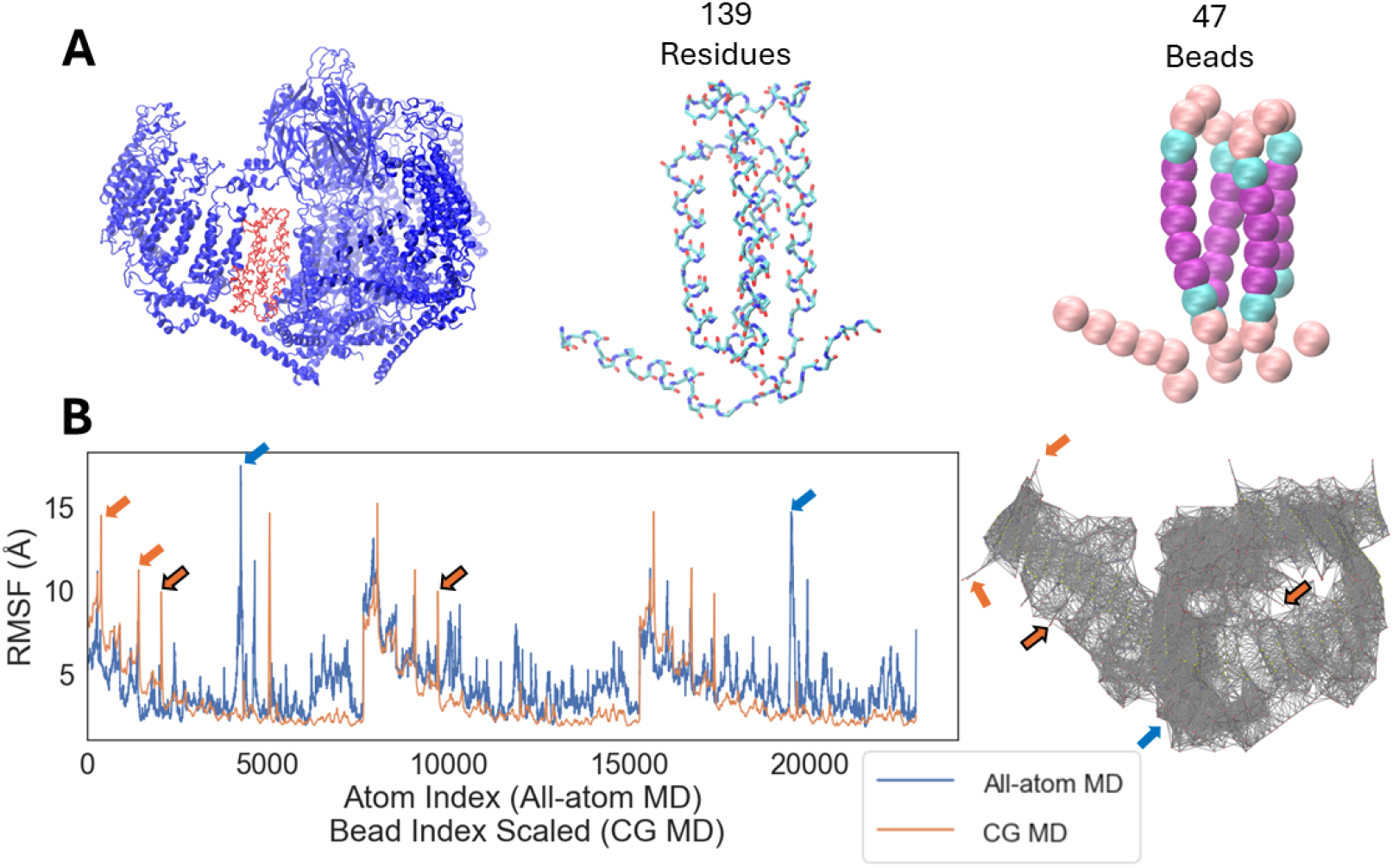
(A) Coarse-graining the structure of PIEZO. The cryogenic electron microscopy structure of full-length PIEZO (PDB ID: 6KG7)^43^ is shown as ribbon diagram in the left panel (blue, with the 139-residue PIEZO repeatA unit in red). Middel panel shows the backbone of repeatA unit as sticks. Right panel shows the corresponding CG-mapped repeatA unit consisting of 47 beads as spheres (bead coloring is consistent with Fig. 1). (B) Root-mean-square fluctuation (RMSF) of protein backbone computed from simulations of all-atom and CG PIEZO. The elastic bonds connecting the CG beads in PIEZO are visualized in the panel on the right. The arrows indicate that the positions of mismatched peaks are due to highly flexible loop regions

### Simulation procedures

#### CG systems

Bilayer geometries were constructed by placing the lipids on an ideal (square) X-Y grid defined by average positions for the equilibrated APL. In simulations containing PIEZO, the protein was positioned into the membrane and overlapping lipids (within *𝓁* = 7.5*Å*) were deleted. Following this, the system underwent a process of minimization and equilibration, during which lipids diffused and filled any voids between the PIEZO arms. The resulting equilibrated system served as a template for subsequent systems.

Simulations were performed with LAMMPS (version 24Mar2022),^59,60^ using the NoséHoover extended Lagrangian formalism for controlling pressure and temperature. The integration time step used in all simulations is *τ* = 50*fs*. The weak coupling times of 100, 000*τ* were used to minimize perturbations of the evolution of the system. The simulations were carried out in both *NPT* (*T* = 300*K, P* = 0*atm*) and *NV T* (*T* = 300*K*) as appropriate to explore the mechanical properties of this new model.

### All-atom simulation

A reference simulation of the PIEZO2 structure resolved by cryogenic electron microscopy (PDB ID: 6KG7)^43^ was prepared using CHARMM-GUI tools.^61–65^ The system was simulated with Gromacs (version 2023.3),^66^ starting with energy minimization for 10 *ps*, equilibration for 250 *ps*, and production for 2 *ns* in vacuum. The temperature (*T* = 303.15*K*) was maintained with a Nosé–Hoover thermostat.

### Analysis of trajectories

We used in-house python scripts relying on packages MDAnalysis, ^67,68^ numpy,^69^ pandas,^70^ to perform all analyses. Visualization and plotting was done using matplotlib,^71^ seaborn,^72^ Microsoft Excel (version 16.84),^73^ VMD (version 1.9.4),^74^ OVITO (version 3.10.3).^75^ APL was estimated as the time-averaged X-Y projected APL based on the equilibrated box dimensions and averaged over three independent replicas. Membrane bending stiffness, or bending modulus, *k*_*C*_, was determined using the fluctuation spectrum method.^76^

### Root-mean-square fluctuation

In order to parameterize the PIEZO ENM model, the per-residue fluctuations was estimated from an all-atom simulation of PIEZO. The protein backbone root-mean-square fluctuations (RMSF) were calculated for both atomistic and CG models in vacuum after aligning the simulated trajectories to their respective initial frame using VMD.^74^

### Lateral pressure

The lateral pressure components along X and Y, *P*_*xx*_ and *P*_*yy*_, respectively, were obtained directly from the stress tensor in LAMMPS at every 5, 000 steps. The X-Y averaged instantaneous lateral pressure is given by (*P*_*xx*_ + *P*_*yy*_)*/*2. A rolling-average procedure was used to denoise the pressure time series. The lateral pressure of the membrane indicates the lateral stress of the membrane with *P* = 0 corresponding to the desirable “stressless” state in the CG ensemble. We remark that the interpretation of pressure in CG simulations is non-trivial^77^ and further that the usual equation for surface tension (*γ* = *L*_*z*_*/*2 × (*P*_*zz*_ − (*P*_*xx*_ + *P*_*yy*_)*/*2), where *L*_*z*_ is the Z box dimension) may not hold, especially considering the significant curvatures manifested in the PIEZO dome and footprint.

### Piezo membrane footprint

The PIEZO footprint was calculated by averaging the position of membrane interfacial beads around the protein. The lateral (2-dimensional) radial distribution function of the interfacial beads was computed based on their X-Y coordinates. The corresponding Z average was computed in bins of 5*Å*.

## Data and code availability

Lipid parameters (original and new), the final CG PIEZO model, example input files, and all analysis scripts are provided on github (https://github.com/charliewen95/LAMMPS-CG-membrane.git).

## Results and discussion

### 4-bead Grime lipid model for describing bilayer membranes

Bilayer membranes formed by the 5- and 4-bead lipid models are attractive for various reasons. First, they (like other resolutions) can have their membrane bending stiffness controlled by a single parameter, the angle potential parameter (BEND), to form a fluid-phase bilayer membrane with mechanical bending stiffness in the range of a few to ∼ 100*k*_*B*_*T* . Furthermore, the 5- and 4-bead resolution are intuitively appealing due to their ability to semi-quantitatively reproduce the correct lateral pressure profile with high-pressure peak in the interfacial region of each leaflet of the membrane.^10,14^ This is an important structural feature, especially when studying transmembrane proteins that cross traverse the membrane. The 4-bead model is preferred over the 5-bead model primarily because of the fewer beads needed to describe the lipid molecule, making the simulation more computationally efficient. Another tangible benefit is that since the 4-bead membrane is less thick than the 5-bead model, it is more responsive to a given change in the angle potential parameter (BEND), which becomes increasingly important as the membrane size gets bigger. Recall that for a given parameter set, the resulting bilayer membrane gets much less stiff when increasing the size of the system (an effect that is due to the pinning at the box boundary). So, the angle parameter will have to be tightened to simulate a bilayer membrane at similar bending stiffness when the box size increases.^10^ Alternatively, the range of the attractive cohesion interaction may be increased.^78^

What is rather unexpected, however, is that the membrane behavior can qualitatively change by inceasing the box size. For instance, we observe that pores, not present in smaller systems, can appear above a certain critical size for a given parameter set. Furthermore, what may be negligible spontaneous curvature in a given simulation box size can give rise to an instability characterized by the membrane bilayers tendency to curl up and buckle when the box size increases. A comphensive characterization to these two effects, and how to deal with each in a pragmatic manner, is presented in the following sections.

### Resolving the scale-dependent limitations

#### Poration due to lipid molecular shape

Using the proposed parameters for the 4-bead Grime lipid model, we can readily realize fluid bilayer membranes with the properties previously reported. ^10^ However, increasing the system size from a box containing 20*k* to 45*k* lipids leads to the formation of pores in the membrane (Fig. 3). For this particular case, we are observing persistent pores that are stable and diffuse around in the bilayer. We note that pores of course can occur in smaller systems as well, but the original parameters were proposed to exclude this undesirable artifact and it was not observed in any of the tested morphologies for smaller systems (*<* 20*k* lipids).

**Figure 3:**
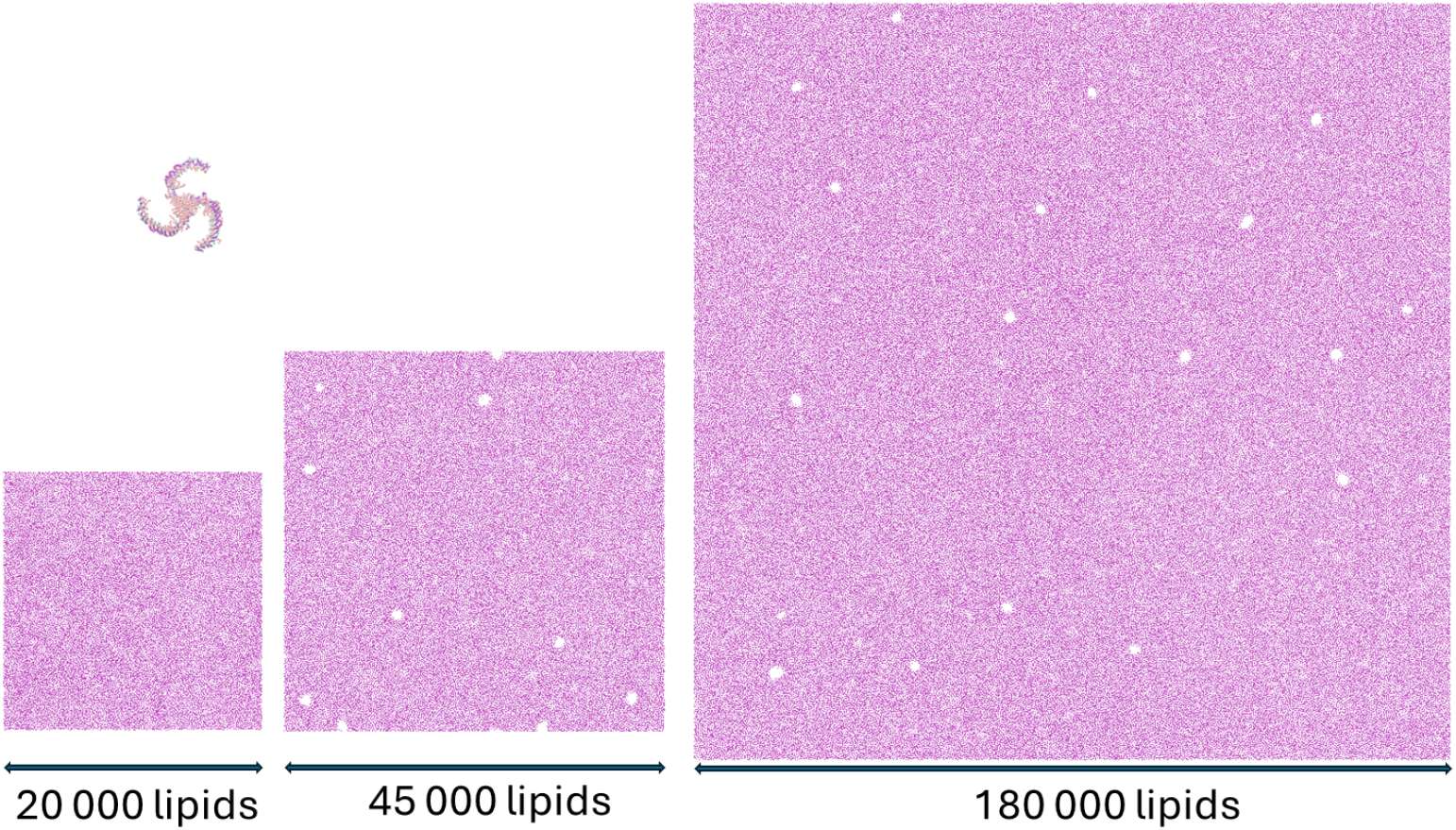
The three membrane sizes simulated are 20*k* lipids (*X* = *Y* ≈ 78.5nm), 45*k* lipids (*X* = *Y* ≈ 115.3nm) and 720*k* lipids (*X* = *Y* ≈ 228.8nm). The poration issue can be seen above the critical size for the reference set of parameters at around or above 45*k* lipids, while completely absent for the system of 20*k* lipids. The size of the PIEZO protein is indicated for comparison.

As has been alluded to previously^79^ and demonstrated to be true for the Grime lipid,^10^ the major characteristic of lipid (or detergent) in determining the assembly morphology is the molecular packing ratio – or how “conical” the molecule is. This immediately suggests that decreasing the size of the head bead in the Grime lipid should help resolve the poration issue, which is indeed what we observe. In fact, all issues of poration, at any size tested, is completely removed by decreasing the size of the lipid head bead from *R*_*HH*_ = 0.75*𝓁* to *R*_*HH*_ ≤ 0.70*𝓁*, where *𝓁* = 7.5 Å is the characteristic length scale (e.g, the radius of the tail beads) in the simulation (Fig. 4).

**Figure 4:**
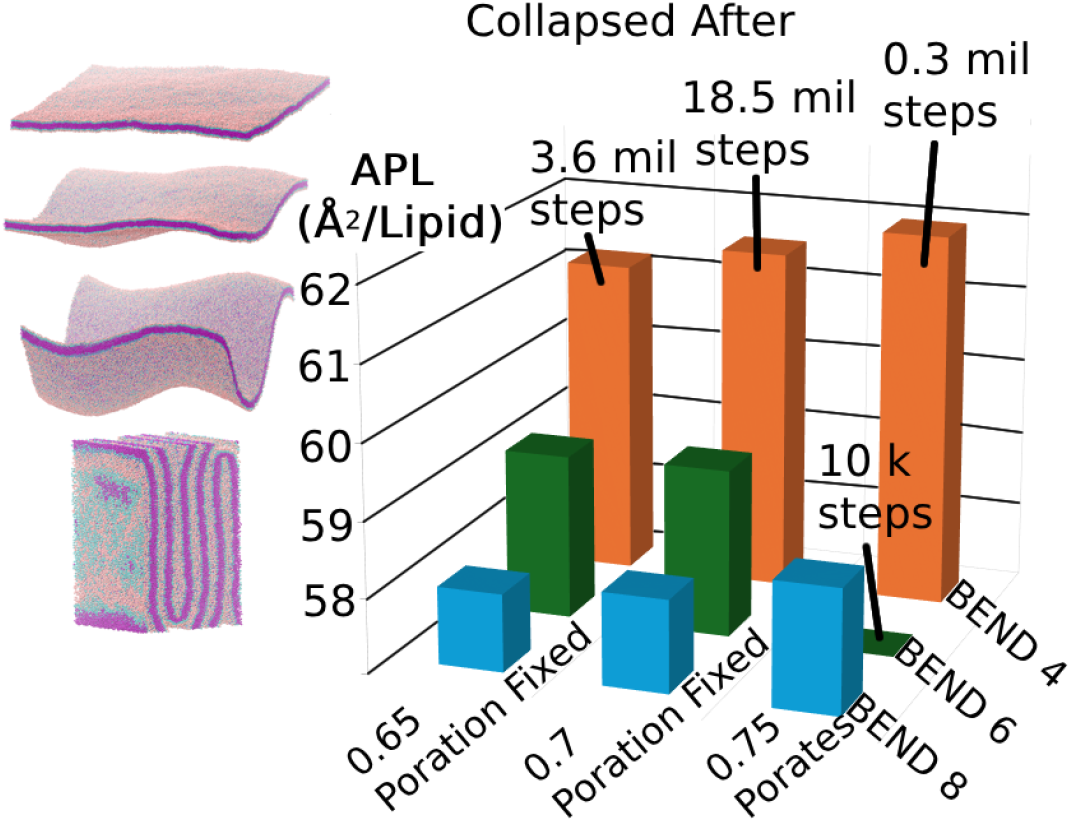
(Left) A series of simulation snapshots show the stages of ever-increasing sinusoidal distortion for a membrane system that buckles. (Right) Exploration of how the head bead size, *R*_*HH*_, and (unit-less) angle parameter, BEND, affect area per lipid (APL), poration and stability. APL values are derived from averaging over three independent replicas. The marked bars correspond to simulations that will collapse in at least one of the three replicas.

### Buckling due to non-vanishing lateral stresses and spontaneous curvature

Upon resolving the issue with poration, we can stably simulate bilayers (without pores) in the *NPT* ensemble for some, but not all, of the desirable range of angle potential parameter (Fig. 4). The stiffer membranes can be simulated with a barostat coupled, but less stiff membranes are often seen to buckle, even if the extrapolated membrane bending modulus, *k*_*C*_, is in a range suggesting stability. A pragmatic approach for this would involve simulating instead the corresponding canonical (*NV T*) ensemble at an appropriate box size.

We note that simulating in the *NV T* as opposed to the *NPT* ensemble usually is not a strong limitation since sampling in the two ensembles would be consistent for large-enough systems.^80,81^ However, for bigger systems (e.g., 180*k* lipids) we tend to see that there is no “good” choice of box size in the sense that the lateral-stress free state (⟨*P*_*xx*_, *P*_*yy*_⟩ ≈ (0, 0)) cannot be achieved. This observation, which at first seems baffling, can be traced to the fact that while the smaller system (∼ 20 to 45*k* lipids) can be simulated in its fluid phase as a (on average) flat bilayer membrane with its characteristic height undulations, increasing the size to e.g. 180*k* lipids causes sinusoidal distortions to persist even while the lateral pressure is non-vanishing and negative. The negative sign of the lateral pressure suggests that the membrane favors contraction, and one might think that the box is simply too large and resizing for a smaller box can resolve the issue. However, this is not the case: decreasing the box size will increase the amplitude of the sinusoidal distortion without ever reaching the state of vanishing lateral stress (Fig. 5). We note that these simulations were performed in the canonical (*NV T*) ensemble so that the box size could be precisely controlled. In all cases for the large system of 180*k* lipids, increasing the box size inevitable leads to increasing (negative) lateral pressure as expected, but decreasing the box size leads to ever-larger amplitude sinusoidal distortions and the persistence of (negative) non-zero lateral pressure (Fig. 5).

**Figure 5:**
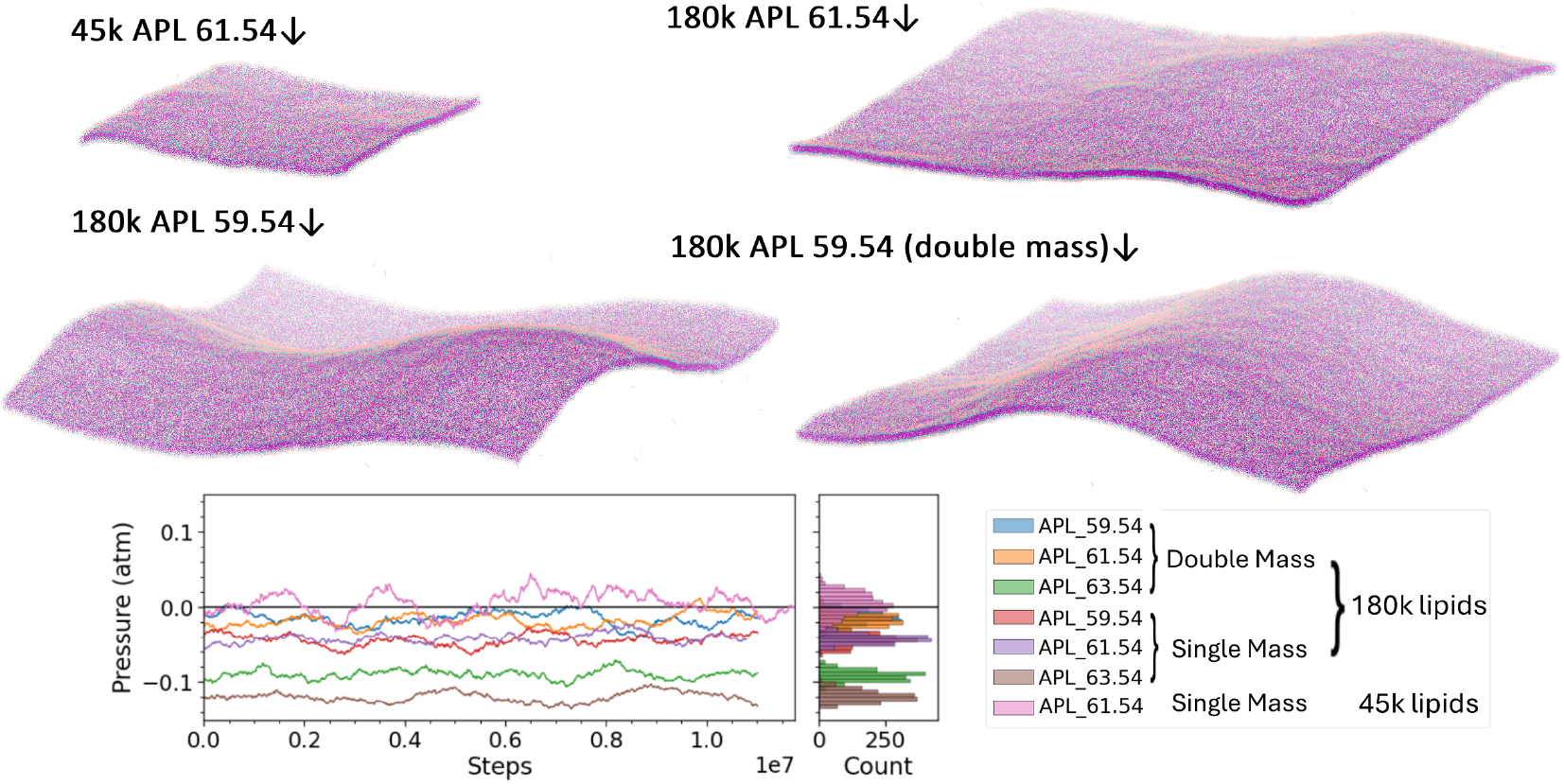
Sinusoidal distortions and non-vanishing lateral pressure in large (180*k* lipids) but not moderately sized (45*k* lipids) bilayers. 45*k* lipids using the pore-fixed parameter set (*R*_*HH*_ = 0.7) and reference-mass beads results in a flat bilayer on the average. For the bilayer with 180*k* lipids, BEND=4 corresponds to a membrane stiffness of *k*_*C*_ = 25*k*_*B*_*T* .

A pragmatic way of increasing stability and reduce the tendency of membrane buckling (in *NPT*) to allow for slightly longer simulations is increase the mass of the beads. Since **F** = *m***a**, this effectively alters the force field and slows the dynamics, though without changing the low-*q* height fluctuation modes of the membrane and hence bending stiffness. However, while simulations with double mass beads were somewhat easier to control, the lateral pressure issue was ultimately not solved using this strategy (Fig. 5).

That vanishing lateral stress is unachievable using the original parameters (or those fixed for poration at large scale) no matter how the box size is altered strongly suggests that the membrane has non-zero spontaneous curvature, *c*_0_ *>* 0. The most poignant demonstration to support this notion is our simulation of a free-standing bilayer disc membrane. Using the original (or “poration-fixed”) parameters, a large enough disc will spontaneously curl up and morphologically transform into its corresponding vesicle (Fig. 6).

**Figure 6:**
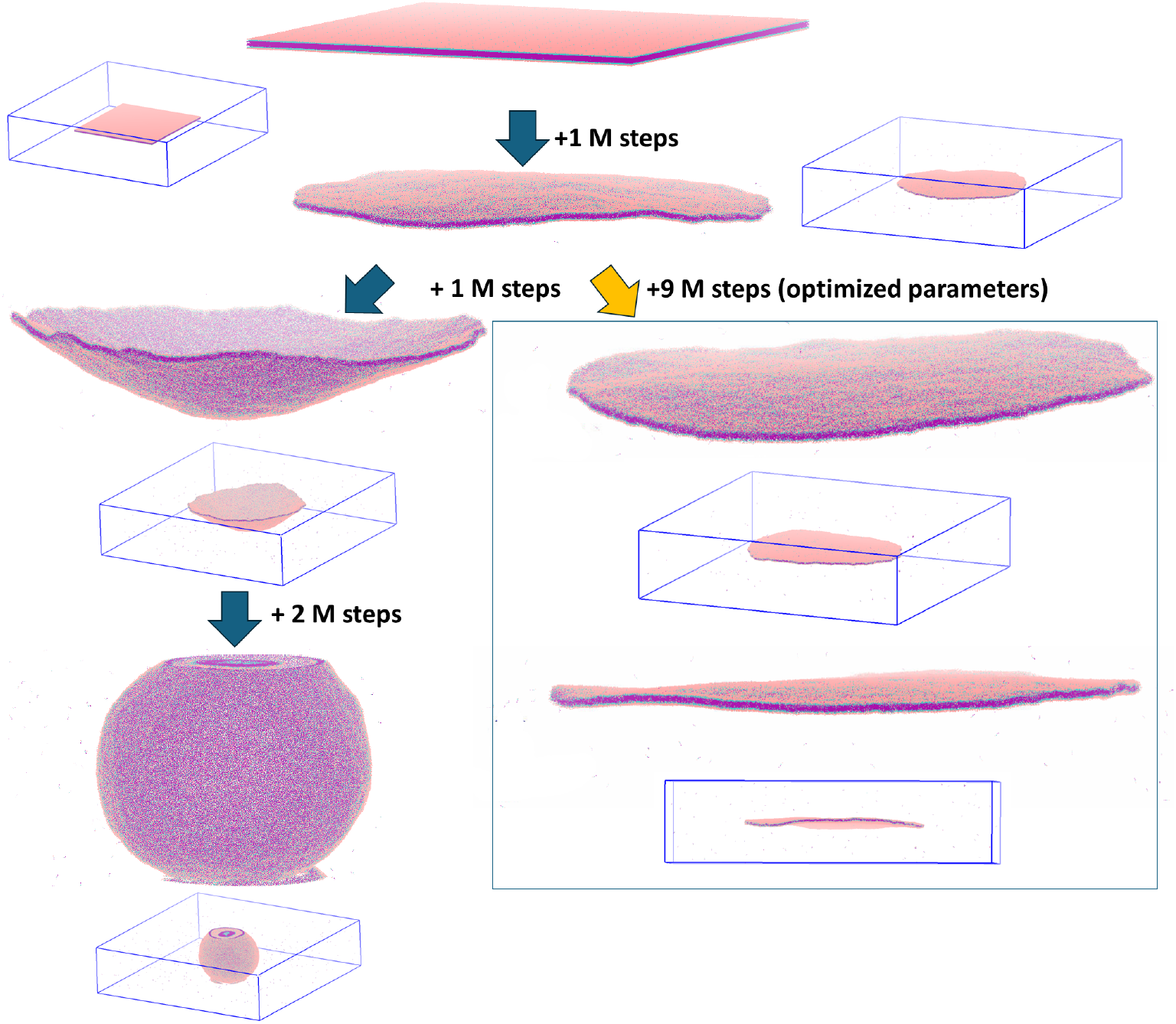
Illustration of the morphological transition from a planar bilayer disc to vesicle caused by the presence of unintentional spontaneous membrane curvature, *c*_0_, caused by the lipid parameters. Lipid positions are initialized in a square grid (top), which relaxes into a disc shape and gradually curls up into a sphere when using the original parameters over 2-4 million MD steps (left). The optimized parameters result in a bilayer disc that remains flat for *>* 10 million MD steps, consistent with *c*_0_ ≈ 0 (right). The smaller panels show the corresponding snapshot in the simulation box (in blue).

Unfortunately, making the head bead, *R*_*HH*_, smaller did nothing to resolve the spontaneous curvature issue. Several other attempts at tweaking the poration-fixed model (*R*_*HH*_ = 0.7*𝓁*), including modifying in addition the angle potential parameter (BEND) or the strength or range of the cohesion (in combination with the head bead-size modification that eliminated poration) were all unsuccessful. A common situation observed in our testing was that there was no usable range where tuning a single lipid parameter would result in a fluid bilayer membrane that was stable (without lipid ejection), without pores, with vanishing lateral stress in its fluid phase over the desirable range of bending stiffness, a few to ∼ 100*k*_*B*_*T* . Lowering the strength of the cohesion rapidly gives rise to stability issues characterized by lipid ejection, and strengthening it gives rise to exclusively too-stiff or even gel-phase states.

### Properties of an optimized 4-bead model

A solution that did prove effective was to solve the poration issue by changing the lipid molecule shape not by resizing the head bead but both the head and interface beads. When this is done in conjunction with a modification to the interface-bead cohesion (increasing the range slightly to facilitate stability), the resulting model will produce a stable fluid bilayer membrane that can be simulated without pores, lateral-stress free, and with vanishing spontaneous curvature over the time and length scales we tested (Fig. 6). We note that the parameter space of the model most certainly allows for other solutions as well and we do not yet have a comprehensive overview of pros and cons of the various options.

The optimized model enables us to conduct simulations with vanishing lateral stress on the average, ⟨*P*_*xx*_, *P*_*yy*_⟩ ≈ (0, 0). However, this does not always make it practical to employ standard barostat algorithms and parameters due to the solvent-free nature of the CG model and increasing amplitude of height fluctuations in large systems. For this reason, we suggest a pragmatic solution whereby the canonical (*NV T*) ensemble is employed, and the initial guess for the box size is estimated from extrapolation of the lipid area-per-lipid, and modified by fine-tuning this value to achieve vanishing lateral stress (Fig. 7A).

**Figure 7:**
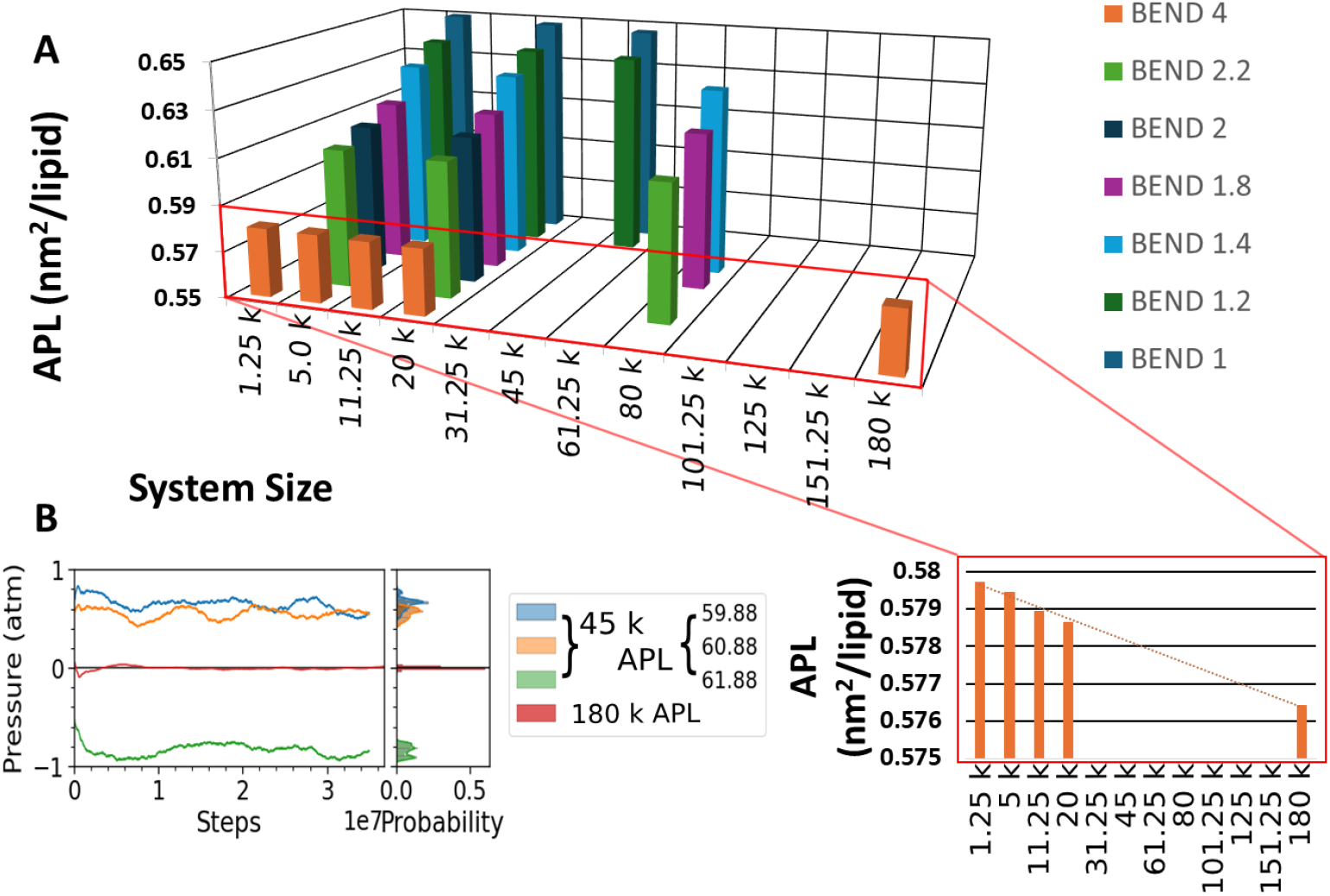
(A). Area per lipid (APL) scales linearly with membrane size across the tested angle parameter BEND values. (B). APL and the resulting lateral pressure ((*P*_*xx*_ + *P*_*yy*_)*/*2) of the optimized 4-bead lipid model with BEND=*X*. The linear trend in APL can be used to extrapolate the appropriate simulation box size for the given setup (BEND value of the lipid and number of lipids) to materialize the lateral stress-less bilayer membrane.

Using this approach, we can realize both negative and positive lateral pressures in small- and medium-sized systems (e.g., 20 − 45*k* lipids) and by using the extrapolation and fine-tuning scheme we can achieve vanishing lateral pressure in the much-larger system of 180*k* lipids (Fig. 7B).

### Large-scale membrane simulation of PIEZO

Using our optimized CG lipid model, we were able to simulate a single PIEZO channel in a stable membrane patch of 115x115*nm*^2^ (Fig.3C). This allows us to investigate the systemic effect of the membrane’s mechanical properties on the PIEZO footprint. Our PIEZO model is based on the cryo-EM structure of full-length PIEZO2 (PDB ID: 6KG7)^43^ solved in detergent, which has an inverted dome shape with an axial height of 9.1 nm and a projected radius of 11 nm.^43^ The shape and size of PIEZO’s footprint depends on the shape of the PIEZO protein, the membrane bending rigidity, *k*_*C*_, and the membrane tension, *γ*.

To probe the effect of membrane rigidity, we first fixed protein conformation and simulated PIEZO in a tensionless bilayer with *k*_*C*_ spanning a physiologically relevant range from 18 to 34*k*_*B*_*T* (Fig.8A). As expected, the footprint size is infinite under zero tension;^44^ hence, the membrane flattens asymptotically up to the boundary of the simulated bilayer. Interestingly, under this condition, the curvature of the membrane decays faster as the membrane becomes more rigid. However, due to the infinite size of the footprint, we cannot entirely exclude the influence of periodic boundary conditions (PBC) on the PIEZO membrane footprint at zero tension.

Next, we kept the fixed protein conformation and membrane bending rigidity *k*_*C*_ of 18 *k*_*B*_*T* . As the membrane being stretched, PIEZO footprint size reduced gradually (Fig.8B). It was found that the membrane lateral pressure remains close to zero up to 14, 200*nm*^2^, beyond which a sudden drop of lateral pressure to -0.5atm was observed (Fig 8C). At the membrane size of 14, 200*nm*^2^, the bilayer becomes nearly flat at 40*nm* from the protein center and 36*nm* from the simulated bilayer boundary. Thus, this system represents PIEZO footprint size under a low resting tension with minimum PBC artifact. If one adopts the relationship between footprint decay length and membrane property predicted from a continuum model^44^ (footprint decay length 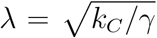), a *k*_*C*_ of 18*k*_*B*_*T* and a *λ* of 29*nm* (calculated as the

**Figure 8:**
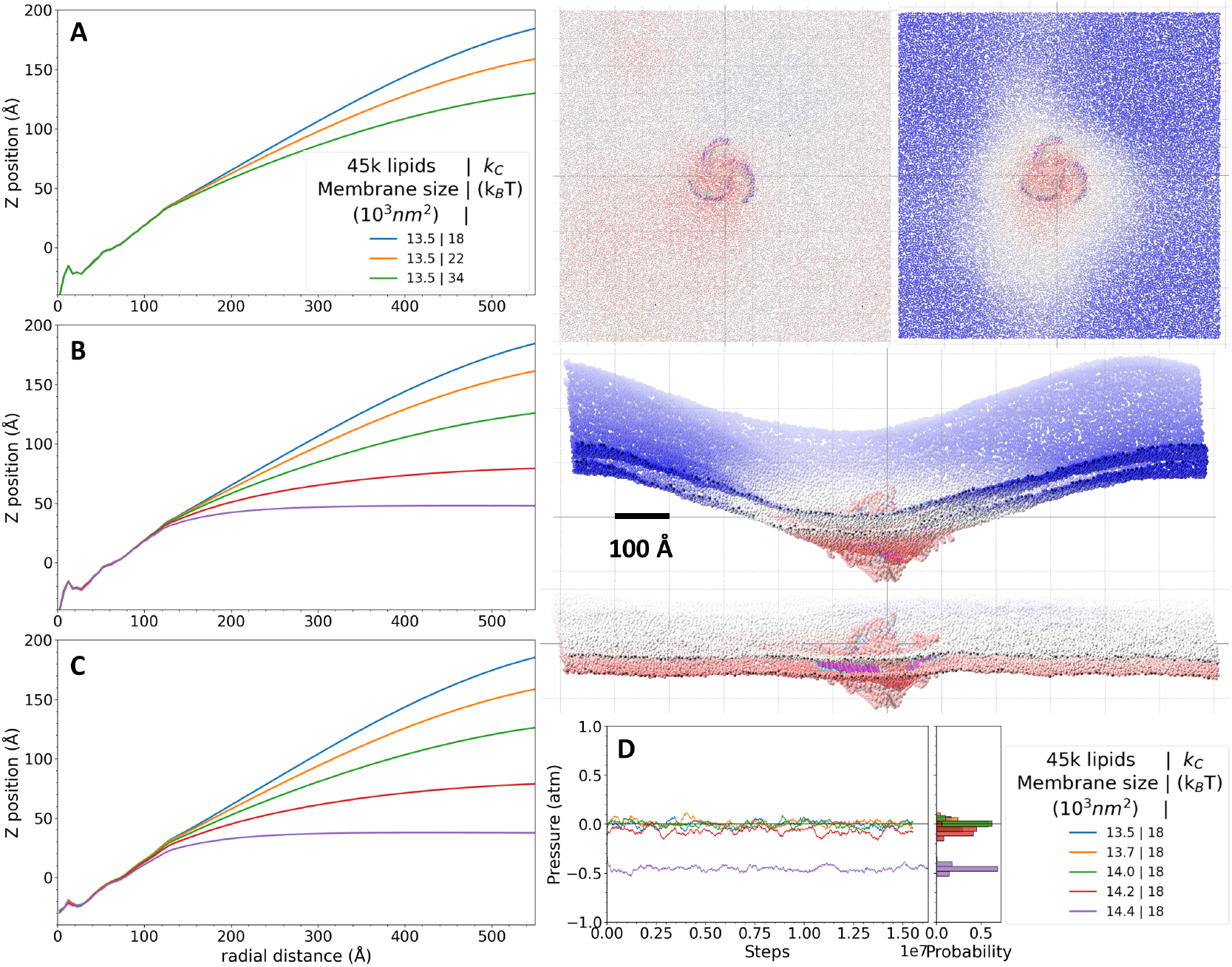
Average membrane Z position as function of radial distance to PIEZO’s center of mass, revealing PIEZO’s membrane footprint in various situations. The simulation box has 45*k* lipids. (A). Protein conformation is fixed and embedded in membranes with increasing bending stiffness, *k*_*C*_ = {18, 22, 34} *k*_*B*_*T* . Protein conformation remains fixed (B) or is allowed to fluctuate (C) upon varying the membrane lateral pressure. (D). (Top Right) Illustration of PIEZO footprint corresponding to panel C with membrane colored by Z-coordinates (along membrane normal) in the absence (membrane size 13.5 x 10^3^*nm*^2^) or presence of lateral stress (membrane size 14.4 x 10^3^*nm*^2^).

distance from the protein center where the membrane flattens minus the projected radius of PIEZO, 40*nm* − 11*nm* = 29*nm*) in our model would yield a low membrane tension of 0.02*k*_*B*_*T/nm*^2^, well within the resting tension of living cell (*γ* = 0.1*k*_*B*_*T/nm*^2^).

To test how our elastic network PIEZO model responds to membrane tension, we conducted the same set of simulations without fixing the protein conformation (Fig.8D). We found that at low resting tension, the PIEZO dome has negligible change, but at lateral pressure of −0.5*atm*, the curvature of the PIEZO dome starts to reduce, consistent with how PIEZO arms sense membrane tension.

## Conclusions

At the micron scale, the cell membrane and its embedded proteins display amazing complexity in their shape and organization. To explore the underlying physical principle driving such phenomena there is a need to simulate the equilibrium ensemble and dynamical behavior of protein diffusion, long-range protein-membrane and protein-protein interactions. The CG model employed here is designed to fill the gap between the elastic continuum model of membrane deformation and atomistic MD approach.

We demonstrate how to systematically resolve scale-dependent issues arising only in systems above a certain critical size, including poration and sinusoidal distortions leading to buckling. While this was done for a specific CG lipid model, we anticipate that the insights gained will be useful for resolving identical problems in similar CG methodologies.

Using our newly optimized CG lipid model, we show that increases in membrane bending rigidity or lateral tension lead to a reduction in the PIEZO dome and membrane footprint. While the extent of the CG protein protein conformational change requires further investigation, ideally in conjunction with atomistic simulations, the response of the PIEZO dome and footprint to membrane rigidity and tension demonstrates the capability of our model in investigating long-range membrane deformation caused by membrane proteins. In particular, the largest membrane patch tested here (230x230 *nm*^2^) will be useful for future simulations of multiple PIEZO channels, which may have intriguing functional consequences if the footprints of multiple PIEZO channels can overlap.

## Acknowledgement

This work was supported by NIH grant 2R01GM130834 to Y.L.L. The authors wish to acknowledge the computational resources provided by WesternU computing cluster and the Advanced Computing Resources at the University of South Florida, as well as Stampede2 and Stampede3 at the Texas Advanced Computing Center through allocation MCB160119 and BIO240033 from the Advanced Cyberinfrastructure Coordination Ecosystem: Services & Support (ACCESS) program,^82^ which is supported by National Science Foundation grants #2138259, #2138286, #2138307, #2137603, and #2138296.

## References

(1) Davies, P. F.; Tripathi, S. C. Mechanical stress mechanisms and the cell. An endothelial paradigm. Circulation Research 1993, 72, 239–245.

(2) Wang, N. Review of cellular mechanotransduction. Journal of Physics D: Applied Physics 2017, 50, 233002.

(3) Delemotte, L.; Luo, Y. Molecular Dynamics. In Handbook of Ion Channels, J. Zheng and M. C. Trudeau (2nd edition); CRC Press, 2023.

(4) Chmiela, S.; Sauceda, H. E.; Müller, K.-R.; Tkatchenko, A. Towards exact molecular dynamics simulations with machine-learned force fields. Nature Communications 2018, 9.

(5) Hollingsworth, S. A.; Dror, R. O. Molecular Dynamics Simulation for All. Neuron 2018, 99, 1129–1143.

(6) Ohkubo, Y. Z.; Madsen, J. J. Uncovering Membrane-Bound Models of Coagulation Factors by Combined Experimental and Computational Approaches. Thrombosis and Haemostasis 2021, 121, 1122–1137.

(7) Lee, S.; Tran, A.; Allsopp, M.; Lim, J. B.; Hénin, J.; Klauda, J. B. CHARMM36 United Atom Chain Model for Lipids and Surfactants. The Journal of Physical Chemistry B 2014, 118, 547–556.

(8) Yang, L.; Tan, C.-h.; Hsieh, M.-J.; Wang, J.; Duan, Y.; Cieplak, P.; Caldwell, J.; Kollman, P. A.; Luo, R. New-Generation Amber United-Atom Force Field. The Journal of Physical Chemistry B 2006, 110, 13166–13176.

(9) Souza, P. C. T. et al. Martini 3: a general purpose force field for coarse-grained molecular dynamics. Nature Methods 2021, 18, 382–388.

(10) Grime, J. M. A.; Madsen, J. J. Efficient Simulation of Tunable Lipid Assemblies Across Scales and Resolutions. arXiv preprint 2019, arXiv:1910.05362.

(11) Grime, J. M. A.; Madsen, J. J. The Grime Coarse-Grained Lipid Model. 2019; https://zenodo.org/record/3479542.

(12) Farago, O. “Water-free” computer model for fluid bilayer membranes. The Journal of Chemical Physics 2003, 119, 596–605.

(13) Kranenburg, M.; Nicolas, J.-P.; Smit, B. Comparison of mesoscopic phospholipid–water models. Phys. Chem. Chem. Phys. 2004, 6, 4142–4151.

(14) Brannigan, G.; Philips, P. F.; Brown, F. L. H. Flexible lipid bilayers in implicit solvent. Physical Review E 2005, 72.

(15) Cooke, I. R.; Kremer, K.; Deserno, M. Tunable generic model for fluid bilayer membranes. Physical Review E 2005, 72.

(16) Lenz, O.; Schmid, F. A simple computer model for liquid lipid bilayers. Journal of Molecular Liquids 2005, 117, 147–152.

(17) Shinoda, W.; DeVane, R.; Klein, M. L. Multi-property fitting and parameterization of a coarse grained model for aqueous surfactants. Molecular Simulation 2007, 33, 27–36.

(18) Orsi, M.; Haubertin, D. Y.; Sanderson, W. E.; Essex, J. W. A Quantitative Coarse-Grain Model for Lipid Bilayers. The Journal of Physical Chemistry B 2007, 112, 802–815.

(19) Revalee, J. D.; Laradji, M.; Sunil Kumar, P. B. Implicit-solvent mesoscale model based on soft-core potentials for self-assembled lipid membranes. The Journal of Chemical Physics 2008, 128.

(20) de Meyer, F.; Smit, B. Effect of cholesterol on the structure of a phospholipid bilayer. Proceedings of the National Academy of Sciences 2009, 106, 3654–3658.

(21) Hömberg, M.; Müller, M. Main phase transition in lipid bilayers: Phase coexistence and line tension in a soft, solvent-free, coarse-grained model. The Journal of Chemical Physics 2010, 132.

(22) Wang, Y.; Sigurdsson, J. K.; Brandt, E.; Atzberger, P. J. Dynamic implicit-solvent coarse-grained models of lipid bilayer membranes: Fluctuating hydrodynamics thermostat. Physical Review E 2013, 88.

(23) Bradley, R.; Radhakrishnan, R. Coarse-Grained Models for Protein-Cell Membrane Interactions. Polymers 2013, 5, 890–936.

(24) Pak, A. J.; Dannenhoffer-Lafage, T.; Madsen, J. J.; Voth, G. A. Systematic Coarse-Grained Lipid Force Fields with Semiexplicit Solvation via Virtual Sites. Journal of Chemical Theory and Computation 2019, 15, 2087–2100.

(25) Dey, S.; Saha, J. SiMPLISTIC: A novel pairwise potential for implicit solvent lipid simulations with single-site models. JCIS Open 2021, 1, 100004.

(26) Ugarte La Torre, D.; Takada, S.; Sugita, Y. Extension of the iSoLF implicit-solvent coarse-grained model for multicomponent lipid bilayers. The Journal of Chemical Physics 2023, 159.

(27) Pak, A. J.; Grime, J. M. A.; Sengupta, P.; Chen, A. K.; Durumeric, A. E. P.; Srivastava, A.; Yeager, M.; Briggs, J. A. G.; Lippincott-Schwartz, J.; Voth, G. A. Immature HIV-1 lattice assembly dynamics are regulated by scaffolding from nucleic acid and the plasma membrane. Proceedings of the National Academy of Sciences 2017, 114.

(28) Flower, T. G.; Takahashi, Y.; Hudait, A.; Rose, K.; Tjahjono, N.; Pak, A. J.; Yokom, A. L.; Liang, X.; Wang, H.-G.; Bouamr, F.; Voth, G. A.; Hurley, J. H. A helical assembly of human ESCRT-I scaffolds reverse-topology membrane scission. Nature Structural and Molecular Biology 2020, 27, 570–580.

(29) Jarin, Z.; Pak, A. J.; Bassereau, P.; Voth, G. A. Lipid-Composition-Mediated Forces Can Stabilize Tubular Assemblies of I-BAR Proteins. Biophysical Journal 2021, 120, 46–54.

(30) Tsai, F.-C.; Henderson, J. M.; Jarin, Z.; Kremneva, E.; Senju, Y.; Pernier, J.; Mikhajlov, O.; Manzi, J.; Kogan, K.; Le Clainche, C.; Voth, G. A.; Lappalainen, P.; Bassereau, P. Activated I-BAR IRSp53 clustering controls the formation of VASP-actin–based membrane protrusions. Science Advances 2022, 8.

(31) Hudait, A.; Hurley, J. H.; Voth, G. A. Dynamics of upstream ESCRT organization at the HIV-1 budding site. Biophysical Journal 2023, 122, 2655–2674.

(32) Pak, A. J.; Yu, A.; Ke, Z.; Briggs, J. A. G.; Voth, G. A. Cooperative multivalent receptor binding promotes exposure of the SARS-CoV-2 fusion machinery core. Nature Communications 2022, 13.

(33) Kim, S.; Li, C.; Farese, R. V.; Walther, T. C.; Voth, G. A. Key Factors Governing Initial Stages of Lipid Droplet Formation. The Journal of Physical Chemistry B 2022, 126, 453–462.

(34) Kim, S.; Chung, J.; Arlt, H.; Pak, A. J.; Farese, R. V.; Walther, T. C.; Voth, G. A. Seipin transmembrane segments critically function in triglyceride nucleation and lipid droplet budding from the membrane. eLife 2022, 11.

(35) Kim, S. Backmapping with Mapping and Isomeric Information. The Journal of Physical Chemistry B 2023, 127, 10488–10497.

(36) Ilias, N.; Richmond, R. V.; Selvarajah, G. T.; Mat Azmi, I. D.; Ajat, M. Structural complexity and physical mechanism of self-assembled lipid as nanocarriers: A review. Asia Pacific Journal of Molecular Biology and Biotechnology 2023, 26–35.

(37) Yu, A.; Pak, A. J.; He, P.; Monje-Galvan, V.; Casalino, L.; Gaieb, Z.; Dommer, A. C.; Amaro, R. E.; Voth, G. A. A multiscale coarse-grained model of the SARS-CoV-2 virion. Biophysical Journal 2021, 120, 1097–1104.

(38) Domingo, M.; Faraudo, J. Effect of surfactants on SARS-CoV-2: Molecular dynamics simulations. The Journal of Chemical Physics 2023, 158.

(39) Hudait, A.; Voth, G. A. HIV-1 capsid shape, orientation, and entropic elasticity regulate translocation into the nuclear pore complex. Proceedings of the National Academy of Sciences 2024, 121.

(40) Kim, H.; Fábián, B.; Hummer, G. Neighbor List Artifacts in Molecular Dynamics Simulations. Journal of Chemical Theory and Computation 2023, 19, 8919–8929.

(41) Coste, B.; Mathur, J.; Schmidt, M.; Earley, T. J.; Ranade, S.; Petrus, M. J.; Dubin, A. E.; Patapoutian, A. Piezo1 and Piezo2 Are Essential Components of Distinct Mechanically Activated Cation Channels. Science 2010, 330, 55–60.

(42) Coste, B.; Xiao, B.; Santos, J. S.; Syeda, R.; Grandl, J.; Spencer, K. S.; Kim, S. E.; Schmidt, M.; Mathur, J.; Dubin, A. E.; Montal, M.; Patapoutian, A. Piezo proteins are pore-forming subunits of mechanically activated channels. Nature 2012, 483, 176–181.

(43) Wang, L.; Zhou, H.; Zhang, M.; Liu, W.; Deng, T.; Zhao, Q.; Li, Y.; Lei, J.; Li, X.; Xiao, B. Structure and mechanogating of the mammalian tactile channel PIEZO2. Nature 2019, 573, 225–229.

(44) Haselwandter, C. A.; MacKinnon, R. Piezo’s membrane footprint and its contribution to mechanosensitivity. eLife 2018, 7.

(45) Tirion, M. M. Large Amplitude Elastic Motions in Proteins from a Single-Parameter, Atomic Analysis. Physical Review Letters 1996, 77, 1905–1908.

(46) Haliloglu, T.; Bahar, I.; Erman, B. Gaussian Dynamics of Folded Proteins. Physical Review Letters 1997, 79, 3090–3093.

(47) Sinitskiy, A. V.; Voth, G. A. Coarse-graining of proteins based on elastic network models. Chemical Physics 2013, 422, 165–174.

(48) Ming, D.; Wall, M. E. Allostery in a Coarse-Grained Model of Protein Dynamics. Physical Review Letters 2005, 95.

(49) Madsen, J. J.; Sinitskiy, A. V.; Li, J.; Voth, G. A. Highly Coarse-Grained Representations of Transmembrane Proteins. Journal of Chemical Theory and Computation 2017, 13, 935–944.

(50) Madsen, J. J.; Grime, J. M. A.; Rossman, J. S.; Voth, G. A. Entropic forces drive clustering and spatial localization of influenza A M2 during viral budding. Proceedings of the National Academy of Sciences 2018, 115.

(51) Madsen, J. J.; Rossman, J. S. In Virus Infected Cells, Subcellular Biochemistry, Vol 106; Vijayakrishnan, S., Jiu, Y., Harris, J. R., Eds.; Springer International Publishing: Cham, 2023; pp 441–459.

(52) Diggins, P.; Liu, C.; Deserno, M.; Potestio, R. Optimal Coarse-Grained Site Selection in Elastic Network Models of Biomolecules. Journal of Chemical Theory and Computation 2018, 15, 648–664.

(53) Foley, T. T.; Kidder, K. M.; Shell, M. S.; Noid, W. G. Exploring the landscape of model representations. Proceedings of the National Academy of Sciences 2020, 117, 24061–24068.

(54) Noid, W. G. Perspective: Advances, Challenges, and Insight for Predictive Coarse-Grained Models. The Journal of Physical Chemistry B 2023, 127, 4174–4207.

(55) Kučerka, N.; Nieh, M.-P.; Katsaras, J. Fluid phase lipid areas and bilayer thicknesses of commonly used phosphatidylcholines as a function of temperature. Biochimica et Biophysica Acta (BBA) - Biomembranes 2011, 1808, 2761–2771.

(56) Huang, J.; MacKerell, A. D. CHARMM36 all-atom additive protein force field: Validation based on comparison to NMR data. Journal of Computational Chemistry 2013, 34, 2135–2145.

(57) Javanainen, M.; Heftberger, P.; Madsen, J. J.; Miettinen, M. S.; Pabst, G.; Ollila, O. H. S. Quantitative Comparison against Experiments Reveals Imperfections in Force Fields’ Descriptions of POPC–Cholesterol Interactions. Journal of Chemical Theory and Computation 2023, 19, 6342–6352.

(58) Kiirikki, A. M. et al. Overlay databank unlocks data-driven analyses of biomolecules for all. Nature Communications 2024, 15.

(59) Plimpton, S. Fast Parallel Algorithms for Short-Range Molecular Dynamics. Journal of Computational Physics 1995, 117, 1–19.

(60) Thompson, A. P.; Aktulga, H. M.; Berger, R.; Bolintineanu, D. S.; Brown, W. M.; Crozier, P. S.; in ‘t Veld, P. J.; Kohlmeyer, A.; Moore, S. G.; Nguyen, T. D.; Shan, R.; Stevens, M. J.; Tranchida, J.; Trott, C.; Plimpton, S. J. LAMMPS - a flexible simulation tool for particle-based materials modeling at the atomic, meso, and continuum scales. Computer Physics Communications 2022, 271, 108171.

(61) Jo, S.; Cheng, X.; Islam, S. M.; Huang, L.; Rui, H.; Zhu, A.; Lee, H. S.; Qi, Y.; Han, W.; Vanommeslaeghe, K.; MacKerell, A. D.; Roux, B.; Im, W. Biomolecular Modelling and Simulations; Elsevier, 2014; p 235–265.

(62) Park, S.-J.; Kern, N.; Brown, T.; Lee, J.; Im, W. CHARMM-GUI PDB Manipulator: Various PDB Structural Modifications for Biomolecular Modeling and Simulation. Journal of Molecular Biology 2023, 435, 167995.

(63) Jo, S.; Kim, T.; Iyer, V. G.; Im, W. CHARMM-GUI: A web-based graphical user interface for CHARMM. Journal of Computational Chemistry 2008, 29, 1859–1865.

(64) Lee, J. et al. CHARMM-GUI Input Generator for NAMD, GROMACS, AMBER, OpenMM, and CHARMM/OpenMM Simulations Using the CHARMM36 Additive Force Field. Journal of Chemical Theory and Computation 2015, 12, 405–413.

(65) Lee, J.; Hitzenberger, M.; Rieger, M.; Kern, N. R.; Zacharias, M.; Im, W. CHARMM-GUI supports the Amber force fields. The Journal of Chemical Physics 2020, 153.

(66) Abraham, M. J.; Murtola, T.; Schulz, R.; Páll, S.; Smith, J. C.; Hess, B.; Lindahl, E. GROMACS: High performance molecular simulations through multi-level parallelism from laptops to supercomputers. SoftwareX 2015, 1–2, 19–25.

(67) Michaud-Agrawal, N.; Denning, E. J.; Woolf, T. B.; Beckstein, O. MDAnalysis: A toolkit for the analysis of molecular dynamics simulations. Journal of Computational Chemistry 2011, 32, 2319–2327.

(68) Gowers, R.; Linke, M.; Barnoud, J.; Reddy, T.; Melo, M.; Seyler, S.; Domański, J.; Dotson, D.; Buchoux, S.; Kenney, I.; Beckstein, O. MDAnalysis: A Python Package for the Rapid Analysis of Molecular Dynamics Simulations. Proceedings of the 15th Python in Science Conference. 2016.

(69) Harris, C. R. et al. Array programming with NumPy. Nature 2020, 585, 357–362.

(70) Wes McKinney Data Structures for Statistical Computing in Python. Proceedings of the 9th Python in Science Conference. 2010; pp 56 – 61.

(71) Hunter, J. D. Matplotlib: A 2D graphics environment. Computing in Science & Engineering 2007, 9, 90–95.

(72) Waskom, M. L. seaborn: statistical data visualization. Journal of Open Source Software 2021, 6, 3021.

(73) Microsoft Corporation Microsoft Excel. https://office.microsoft.com/excel.

(74) Humphrey, W.; Dalke, A.; Schulten, K. VMD: Visual molecular dynamics. Journal of Molecular Graphics 1996, 14, 33–38.

(75) Stukowski, A. Visualization and analysis of atomistic simulation data with OVITO-the Open Visualization Tool. MODELLING AND SIMULATION IN MATERIALS SCIENCE AND ENGINEERING 2010, 18.

(76) Brandt, E. G.; Braun, A. R.; Sachs, J. N.; Nagle, J. F.; Edholm, O. Interpretation of Fluctuation Spectra in Lipid Bilayer Simulations. Biophysical Journal 2011, 100, 2104–2111.

(77) Wagner, J. W.; Dama, J. F.; Durumeric, A. E. P.; Voth, G. A. On the representability problem and the physical meaning of coarse-grained models. The Journal of Chemical Physics 2016, 145.

(78) Cooke, I. R.; Deserno, M. Solvent-free model for self-assembling fluid bilayer membranes: Stabilization of the fluid phase based on broad attractive tail potentials. The Journal of Chemical Physics 2005, 123.

(79) Israelachvili, J. N.; Mitchell, D. J.; Ninham, B. W. Theory of self-assembly of hydrocarbon amphiphiles into micelles and bilayers. Journal of the Chemical Society, Faraday Transactions 2 1976, 72, 1525.

(80) Allen, M. P.; Tildesley, D. J. Computer Simulation of Liquids; Oxford University Press Oxford, 2017; p 46–94.

(81) Tan, S. J.; Prasetyo, L.; Zeng, Y.; Do, D.; Nicholson, D. On the consistency of NVT, NPT, µVT and Gibbs ensembles in the framework of kinetic Monte Carlo – Fluid phase equilibria and adsorption of pure component systems. Chemical Engineering Journal 2017, 316, 243–254.

(82) Boerner, T. J.; Deems, S.; Furlani, T. R.; Knuth, S. L.; Towns, J. ACCESS: Advancing Innovation: NSF’s Advanced Cyberinfrastructure Coordination Ecosystem: Services and Support. Practice and Experience in Advanced Research Computing. 2023.

